# Simultaneous assessment of P50, MMN, ERN and P300 event-related potentials among patients with Schizophrenia – an exploratory study

**DOI:** 10.1101/837815

**Authors:** Arun Sasidharan, Ajay Kumar Nair, Vrinda Marigowda, Ammu Lukose, John P John, Bindu M Kutty

## Abstract

**Purpose:** P50 suppression (sensory gating or inhibition), MMN (mismatch negativity; bottom-up detection of change), ERN (error related negativity; conflict monitoring) and P300 (attention allocation and memory updating for salient events) are event related potentials (ERPs) widely reported to show abnormal cognitive functioning among patients with schizophrenia. In real-life scenarios the brain processing underlying these ERPs occur simultaneously, and yet prior ERP studies have evaluated them in isolation. The current study uses a novel paradigm that can examine these multiple ERPs simultaneously, and explore if the reported ERP deficits would hold true during a more realistic setting.

**Method:** Data from 21 patients with schizophrenia and 25 age- and gender-matched healthy controls were used, who underwent ERP recordings during the Assessing Neurocognition via Gamified Experimental Logic (ANGEL) paradigm. This is a gamified visual odd-ball paradigm that generates P300, the error responses generate ERN, and paired-tone audio distractors generate P50 and MMN. Peak-peak amplitude, mean amplitude and area-under-curve measures of ERP were measured at electrodes reflecting best morphology.

**Results:** Though patients showed apparent ERP morphology differences relative to the controls, the standard ERP measures were comparable between groups, except for reduced ERN among patients. Interestingly, significant group differences were seen in N1-P2 complex suppression, despite comparable P50 suppression.

**Contribution of the research:** The current study is the first to report multiple ERP component measures simultaneously evoked among patients with schizophrenia, and shows greater signs for impaired prediction mechanism. The findings of the study would provide a more ecologically valid evaluation of ERP-based cognitive functioning, and need to be replicated in a larger sample as well as other mental disorders.

## Introduction

Schizophrenia is a debilitating psychiatric illness that with a global incidence of 1% of general population. It is widely considered as a ‘disorder of thinking’, resulting in impaired sense of reality, altered behaviour and deranged cognitive abilities. Currently the diagnosis of schizophrenia is made by a psychiatrist/psychologist based on verbal descriptions and diagnostic scales obtained during structured interviews with patient and their care-takers. There have been several attempts to find biomarkers that rely on the underlying brain function abnormality for the diagnosis of this complex disorder. One such attempt is to use abnormalities in event related potentials (ERP) as a proxy measure of cognitive dysfunctions.

ERPs are waveforms representing consistent changes in an individual’s EEG data associated with repeatedly presenting a stimulus pattern in particular conditions. Traditionally, each ERP has been thought to reflect a cognitive function, but recent studies suggest that such one to one associations may have limitations. P50 suppression (sensory gating or inhibition), MMN (mismatch negativity; bottom-up detection of change), ERN (error related negativity; conflict monitoring) and P300 (attention allocation and memory updating for salient events) are ERPs widely reported to show abnormal cognitive functioning among patients with schizophrenia. One possible explanation that links the above abnormalities and schizophrenia pathophysiology is that the altered sense of reality experienced by schizophrenia patients has been proposed to be due to a faulty ‘prediction mechanism’. In this regard, sensory gating, change detection, conflict monitoring and attention allocations can be considered to be part of brain’s predictive process, which may be deranged in schizophrenia patients. In a natural setting the above brain processes occur simultaneously. However, the ERP deficits reported among schizophrenia patients were evaluated in isolation. Therefore, the currently available ERP findings may not reflect the true cognitive abnormalities in schizophrenia patients. Some of the new initiatives in mental disorders such as Research Domain Criteria (RDoC) also emphasises on the need for clinically practicable and yet reliable assessments for providing reliable biomarkers for cognitive functioning(1). To address this issue, Kappenman and Luck recently came up with an idea of simultaneously acquiring multiple ERPs as manipulation of orthogonal neural systems together (MONSTER)(2). Our lab had improvised the concept by adding more ERPs to capture greater multi-dimensional brain functional data and also gamifying the paradigm to improve subject involvement in the task (Assessing Neurocognition via Gamified Experimental Logic or ANGEL paradigm)(3).

Accordingly, the current study used the ANGEL paradigm to simultaneously assess P50 suppression, MMN, ERN and P300 ERPs among schizophrenia patients, and explore if the previously reported deficits hold true during a more realistic setting. We hypothesised that the ERP deficits in schizophrenia patients would be less pronounced or absent when assessed simultaneously.

## Method

The study uses ERP data that were part of a sleep-ERP-fMRI study carried out at the National Institute of Mental Health and Neurosciences (NIMHANS), Bangalore, India, with approval from the NIMHANS Ethics Committee, thus conforming to the ethical standards laid down in the 1964 Declaration of Helskinki. Written informed consent was obtained from all participants (and their legally qualified representatives in the case of patients with schizophrenia) prior to enrolling them into the study.

### Participants

The study samples comprised 25 healthy controls (CNT; age: 20–42 years, mean 29.05 years; 12 males), who were recruited by word of mouth, and 23 patients with schizophrenia (SCZ; age: 19–44 years, mean 27.71 years, 14 males), recruited from the out-patient department of NIMHANS by purposive sampling. The samples were matched for age and gender. The socio-demographic and clinical characteristics of the samples are given in Error! Reference source not found.. Only right-handed participants (as determined by Edinburgh’s inventory) in the age group 16 to 50 were included in the study. The diagnosis of schizophrenia was arrived at using criteria from DSM–IV(4) based on the consensus of a research psychiatrist who conducted a semi-structured interview and a trained research psychologist who used the Mini International Neuropsychiatric Interview for DSM-IV (MINI-Plus)(5). The medication status of the participants with schizophrenia are given in Table 1. In case of individuals in the control group, the presence of any medical/psychiatric/neurological condition requiring continuous medications, current psychotropic use and history of psychiatric illness in first-degree relatives were ruled out by an unstructured clinical interview.

**Table 1:** Profile of subjects whose data were used in the study.

| **Parameter** | **SCZ (n=21)** | **CNT (n=25)** | **Statistics** |
| --- | --- | --- | --- |
| **Age** | 27.71 ± 7.21<br>(19.0-44.0) | 29.00 ± 4.96<br>(20.0-42.0) | t statistic = -0.687;<br>p-value = 0.496;<br>CI = -4.8 to 2.4 |
| **Education (years)** | 12.45 ± 3.61<br>(3.0-18.0) | 18.92 ± 2.24<br>(13.0-24.0) | t statistic = -6.974;<br>p-value = 0.000;<br>CI = -8.3 to -4.8 |
| **Age of onset of illness (years)** | 24.76 ± 6.99<br>(17.0-44.0) |  |  |
| **Duration of illness (months)** | 36.48 ± 31.05<br>(3.0-120.0) |  |  |
| **Treatment Status** | Regular = 12 (57.1%)<br>Drug naive = 7 (33.3%)<br>Drug free = 2 (9.5%) |  |  |
| Data expressed as mean ± SD (range). Statistical analysis done using permutation-based t-test and bootstrapping-based confidence interval of group difference, both with 10000 re-sampling with significance at p>0.05. SCZ – patients with schizophrenia; CNT – healthy controls; CI – 95% confidence interval of the mean group difference. |  |  |  |

### Task design and presentation

The simultaneous assessment of multiple ERPs in a single session was made possible using a novel game-based audio-visual task named ‘ANGEL’ (Assessing Neurocognition via Gamified Experimental Logic; developed and validated in our laboratory and described in (3)). Briefly, the subject watches a pair of two-tone images (one a known figure and the other a checker pattern) presented on either sides of a screen, and has to press a left or right button based on a set of simple rules for the known figure (that changes with game level). There are three game levels with increasing complexity (Level 1: chance to learn passively; Level 2: additional distraction while responding; Level 3: change to tougher rules), that keep the subjects engaged and allow to assess various interaction effects. Instructions and practice sessions are provided at the start of each game level. The known figure could be from a set of degraded human faces (Mooney Face), illusory shapes (Kanisza triangle) or their scrambled counterparts, all used to evoke gestalt perception(6). These are sequenced into 25 trial blocks such that each block was a visual oddball (to study P300 ERP), with one type of image presented frequently (80% probability or 20 trials) and two other image types were presented rarely (10% probability or 2/3 trials each). Within a block, the frequent image always come on one side, whereas the rare images may come on either sides. There were 8 blocks, each with a different image combination (4 image types and two sides), presented in a pseudo-random sequence. Each trial is 1200ms to 1900ms long. Subjects have up to 700ms to respond to the stimuli (using index fingers of either hands) after which it is considered a missed response. Frequent performance feedbacks are given (every 2 blocks or 50 trials) to keep the subjects engaged and generate errors while trying to improve performance (‘game-like’ effect). Each game level is about 15 minutes in duration and the overall time for EEG-ERP acquisition is about 1.5 hours including time for scalp preparation and briefing the subject. Paired auditory tones (15ms long; 500ms apart) were presented as distractors every third trial (with pseudorandom onset time) during the task, which help study P50 suppression. The tones were either standard (1000Hz; 80% probability) or deviant (1500Hz; 20% probability); this passive odd ball context allowed MMN assessment for the first tone among the pairs. Eprime 2.0 stimulus presentation software (Psychology Software Tools, Inc., Sharpsburg, PA, USA) was used for presenting the paradigm, synchronised with the EEG acquisition. The subjects sat comfortably in a chair with armrest in front of a 41cm × 26cm LCD monitor (Acer, New Taipei, Taiwan) set 96 cm away. Auditory stimuli were presented at 75dB via pneumatic earphones. Subjects used a two button wireless mouse (Logitech, Switzerland) for responding with the left or right index fingers as appropriate.

### EEG recording

All recordings were done in a sound attenuated chamber in the human cognitive research lab, NIMHANS, with ambient temperature maintained at 25°C. The study was carried out either in the afternoons (2-6pm) or mornings (8-12am), counterbalanced between the subjects. EEG was acquired using a 70 channel Neuroscan SynAmps2 acquisition system (Compumedics, Charlotte, USA), digitized with a resolution of 24 bits and sampling rate of 1000Hz, a high pass filter of 0.1-100Hz, and no notch filters. Neuroscan’s gel based sintered-silver electrode caps (Quik caps, Neuroscan-Compumedics, Charlotte, USA) with 64 monopolar electrodes for EEG, and 2 sets of bipolar electrodes for electro-oculogram (EOG) were used. EEG electrode positions were in accordance with the 10-10 international system for electrode placement(7–9). Reference electrode was between Cz and CPz, and ground between Fz and FPz. Horizontal EOG electrodes were placed 2cm away from the outer canthi of both the eyes, and vertical EOG, electrodes were placed 2cm above and below the left eye. Disposable syringes (5ml) with 14G blunt needles were used for applying gel. Impedance for all electrodes was maintained between 5 to 10 KΩ.

### ERP analysis

All pre-processing was performed offline using EEGLAB v13 toolbox(10) implemented in MATLAB 2013a. EEG drift artefacts removed using a 0.25-0.75Hz transition band high-pass filter, bad channels were rejected using an automated correlation based routine, and high amplitude stereotypical (e.g., eye blinks) and non-stereotypical (e.g., movement) artefacts removed using an artefact subspace reconstruction (ASR) algorithm(11); all implemented using a plugin in EEGLAB. ASR relies on a sliding window principal component analysis (PCA) to statistically compare with a reference clean portion of the EEG data and then linearly reconstruct deviant portions of the data (‘artefact subspaces’) based on the correlation structure observed in the reference clean data(11). We used a threshold of 5 standard deviations for ASR. Deleted bad channels were interpolated using spherical spline interpolation and re-referenced to all channel average. Further analysis were done using ERPLAB plugin of EEGLAB(12). The preprocessed EEG data were segmented into 2500ms epochs (1000ms pre-stimulus data) time-locked to event-markers corresponding to the various ERP conditions (frequent and rare images for P300; first and second tones of the standard tone-pairs for P50 suppression; first tones of standard and deviant tone-pairs for MMN; correct and incorrect responses for ERN). Baseline correction for P50 and MMN analysis was −50ms to 0ms, whereas for the rest of the ERP it was −200ms to 0ms. The epochs were then averaged to generate ERPs of different conditions for each subject. The electrodes and time windows for evaluation of ERPs were chosen based on those reported in previous studies and further verifying them on the respective grand average ERPs from all participant data (irrespective of groups). Accordingly, the electrodes (two of the best electrodes), time-windows and measurement parameters chosen for each of the ERPs are shown in Table 1.

### Statistical analysis

All statistical tests for EEG data were implemented using statistical functions implemented in MATLAB 2013a software. Statistical comparison of ERP parameters were computed by non-parametric permutation based t-test using 10000 iterations, and 95% confidence interval of the mean difference were obtained using bootstrapping statistics on 10000 re-samples. To improve the validity of the above two ‘robust’ measures, statistical significance between groups were made based on a consensus between permutation and bootstrapping statistics, with p<0.05.

## Results

After rejecting 2 patient data having less number of epochs, we could conduct ERP analysis on data from 46 subjects (21 patients and 25 controls). Except for correct/incorrect conditions, number of epochs were comparable between the groups (Table 2). The results are summarized in Table 3. As no apparent latency differences were observed in the grand averages, these were not further evaluated.

**Table 2:**
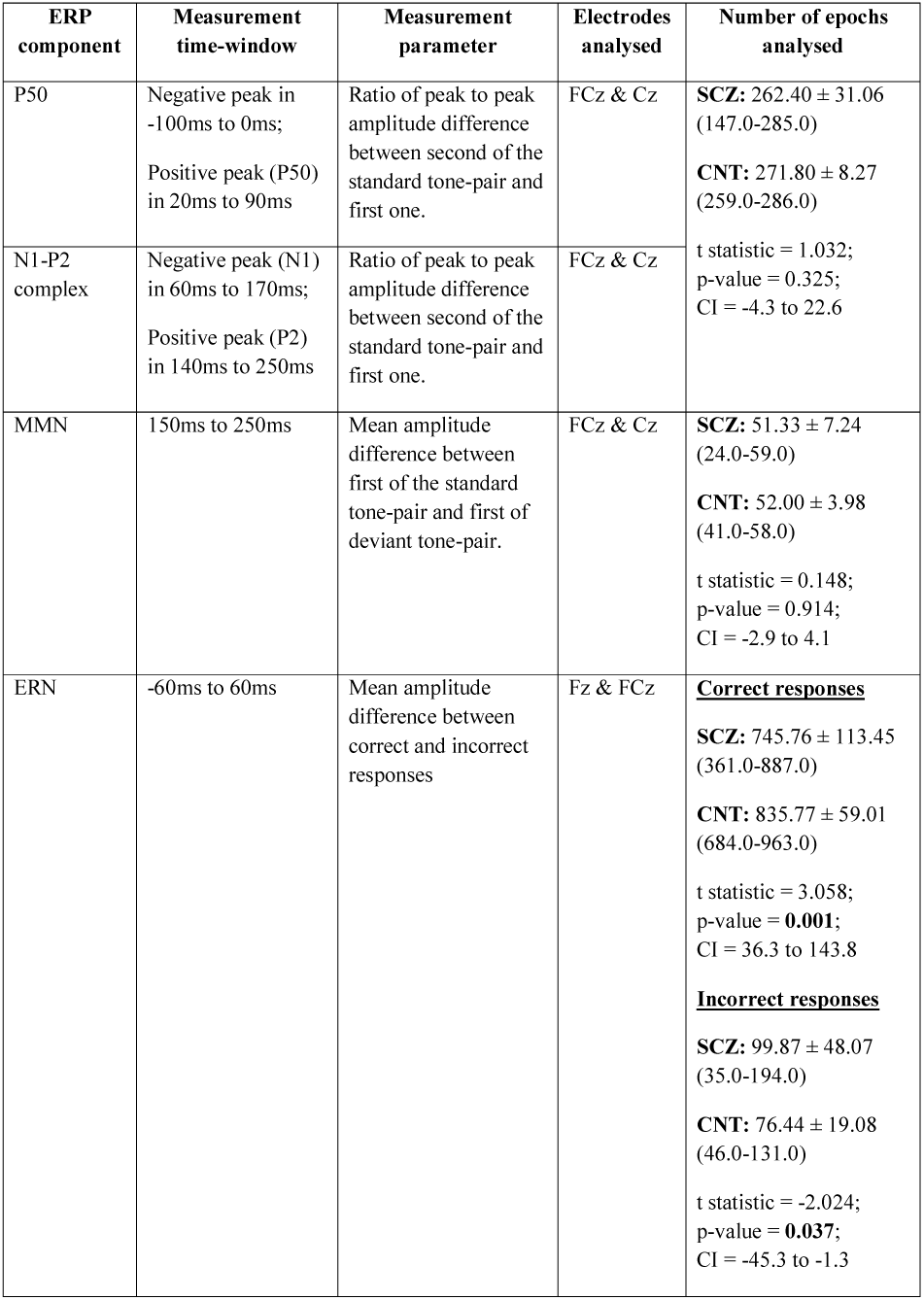
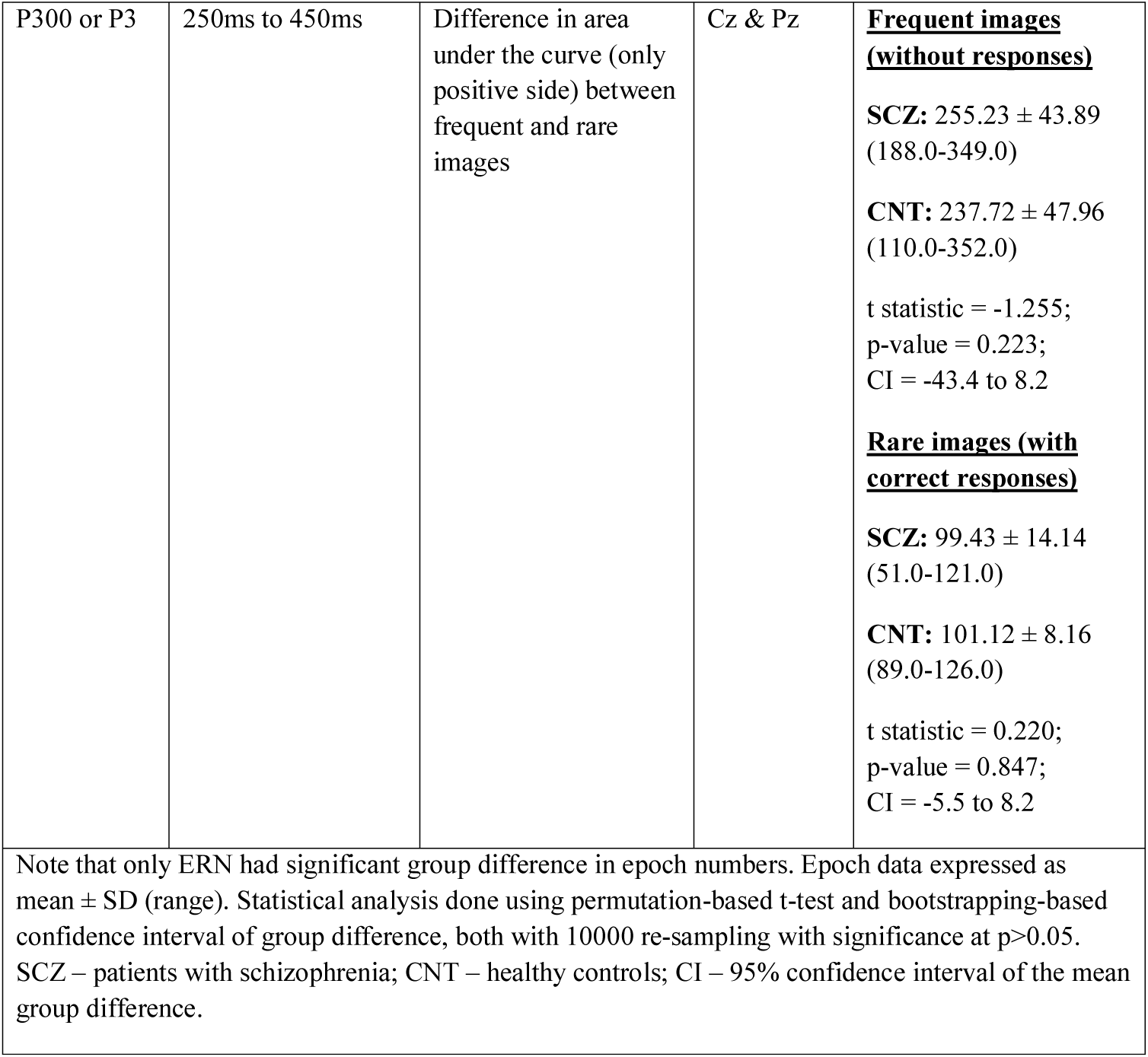
ERP measurement criteria used in the study.

**Table 3:** Results of ERP analysis.

| ERP measure | SCZ (n=21) | CNT (n=25) | Statistics |
| --- | --- | --- | --- |
| P50 suppression | 0.91 ± 0.52<br>(0.3-2.5) | 0.92 ± 0.70<br>(0.2-3.3) | t statistic = -0.024;<br>p-value = 0.972;<br>CI = -0.4 to 0.4 |
| N1-P2 suppression | 0.62 ± 0.31 (0.3-1.6) | 0.46 ± 0.19<br>(0.2-1.0) | t statistic = -1.938;<br>**p-value = 0.049;**<br>**CI = -0.3 to -0.0** |
| MMN | 0.24 ± 0.64<br>(-1.1-1.9) | 0.50 ± 0.66<br>(-0.6-2.2) | t statistic = 1.320;<br>p-value = 0.207;<br>CI = -0.1 to 0.6 |
| ERN | 0.13 ± 1.03<br>(-2.1-3.0) | 1.23 ± 0.87<br>(-0.7-2.6) | t statistic = 3.886;<br>**p-value = 0.000;**<br>**CI = 0.6 to 1.7** |
| P3 | -0.11 ± 0.15<br>(-0.5-0.1) | -0.17 ± 0.19<br>(-0.7-0.1) | t statistic = -0.966;<br>p-value = 0.348;<br>CI = -0.1 to 0.0 |
| Data expressed as mean ± SD (range). Statistical analysis done using permutation-based t-test and bootstrapping-based confidence interval of group difference, both with 10000 re-sampling with significance at p>0.05. Note that N1-P2 suppression and ERN showed significant reduction in patients. SCZ – patients with schizophrenia; CNT – healthy controls; CI – 95% confidence interval of the mean group difference. |  |  |  |

### P50 suppression (Fig. 1)

Despite the apparent reduction of the P50 waveforms among SCZ patients, P50 suppression for the paired tones was comparable between the groups (t statistic = −0.024; p-value = 0.972; CI = – 0.4 to 0.4). But the N1-P2 complex showed significantly lower suppression among the SCZ patients (t statistic = −1.938; p-value = 0.049; CI = −0.3 to −0.0).

**Fig. 1:**
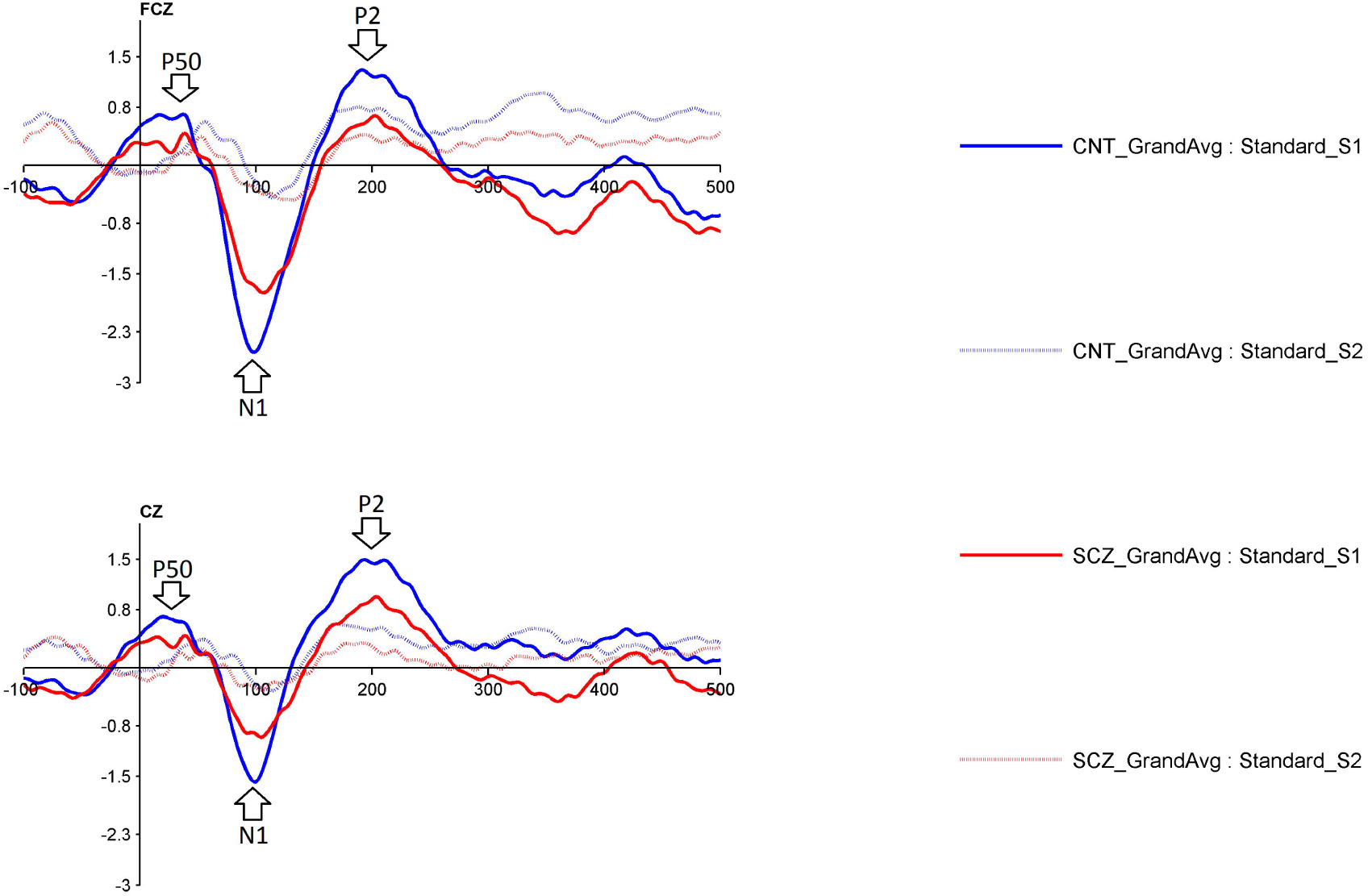
Grand average ERPs showing P50 and N1-P2 suppression in patients with schizophrenia (red) and healthy controls (blue). First and second tones of the standard tone-pairs were the time-locking stimuli. Arrows denote the ERP component studied.

### MMN (Fig. 2)

MMN between standard and deviant tones was comparable between the groups (t statistic = 1.320; p-value = 0.207; CI = −0.1 to 0.6).

**Fig. 2:**
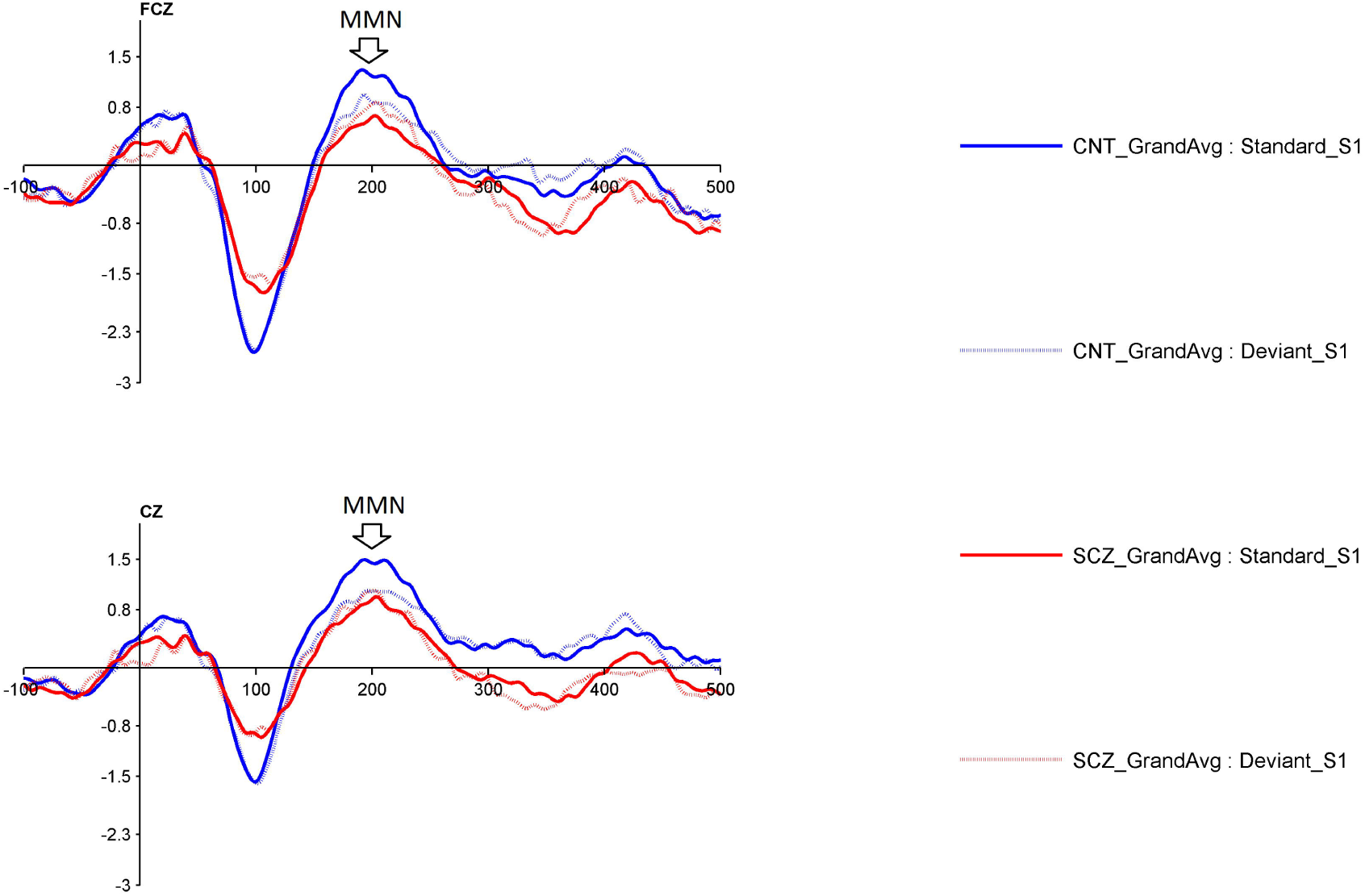
Grand average ERPs showing MMN in patients with schizophrenia (red) and healthy controls (blue). First tones of the standard and deviant tone-pairs were the time-locking stimuli. Arrows denote the ERP component studied.

### ERN (Fig. 3)

SCZ group showed significantly lower ERN between correct and incorrect responses (t statistic = 3.886; p-value = 0.000; CI = 0.6 to 1.7).

**Fig. 3:**
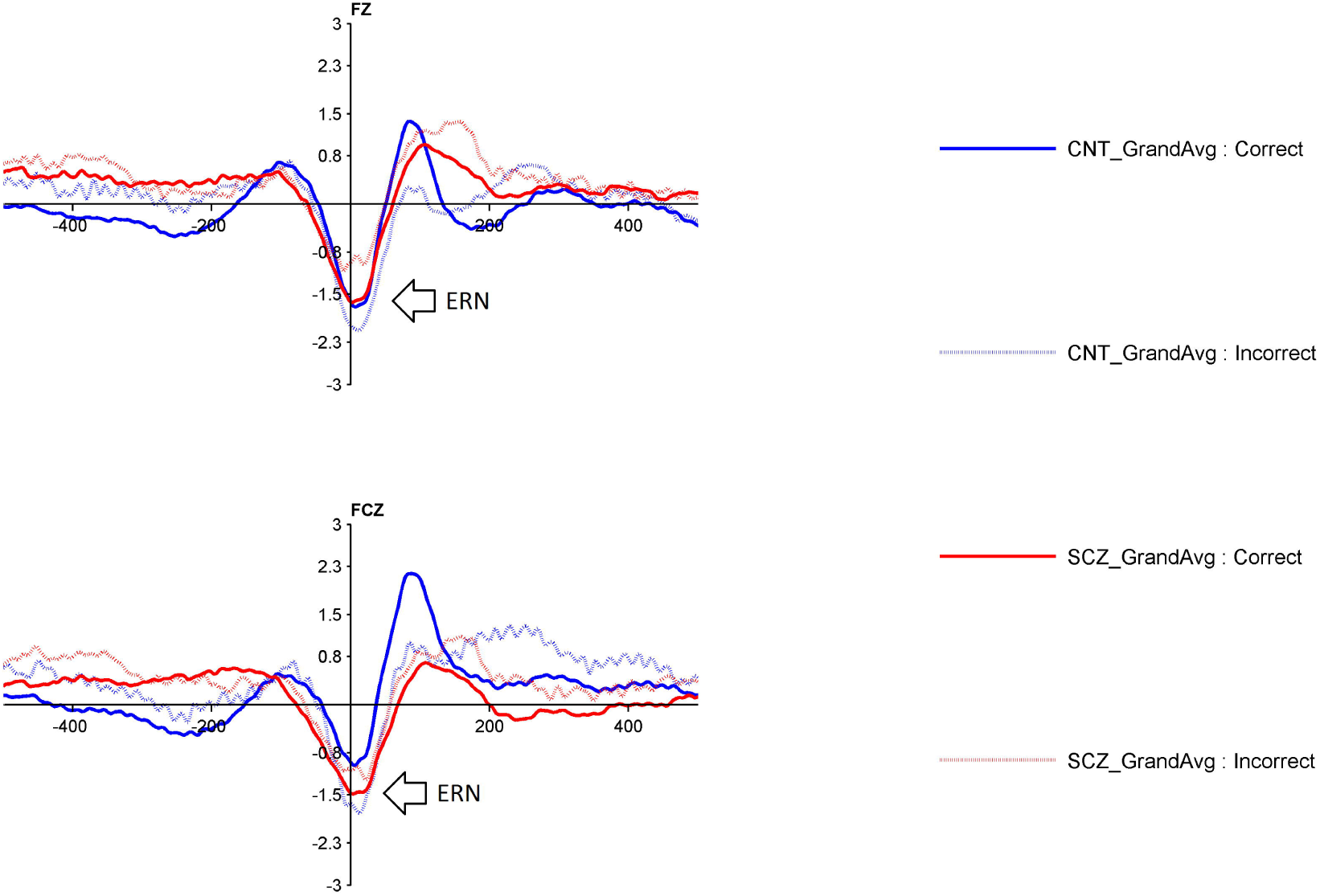
Grand average ERPs showing ERN in patients with schizophrenia (red) and healthy controls (blue). Correct and incorrect responses during the task were the time-locking stimuli. Arrows denote the ERP component studied.

### P300 (Fig. 4)

Though the ERP waveforms were smaller among SCZ patients, both groups showed similar increase in P300 or P3 component during rare compared to frequent visual stimuli (t statistic = – 0.966; p-value = 0.348; CI = −0.1 to 0.0).

**Fig. 4:**
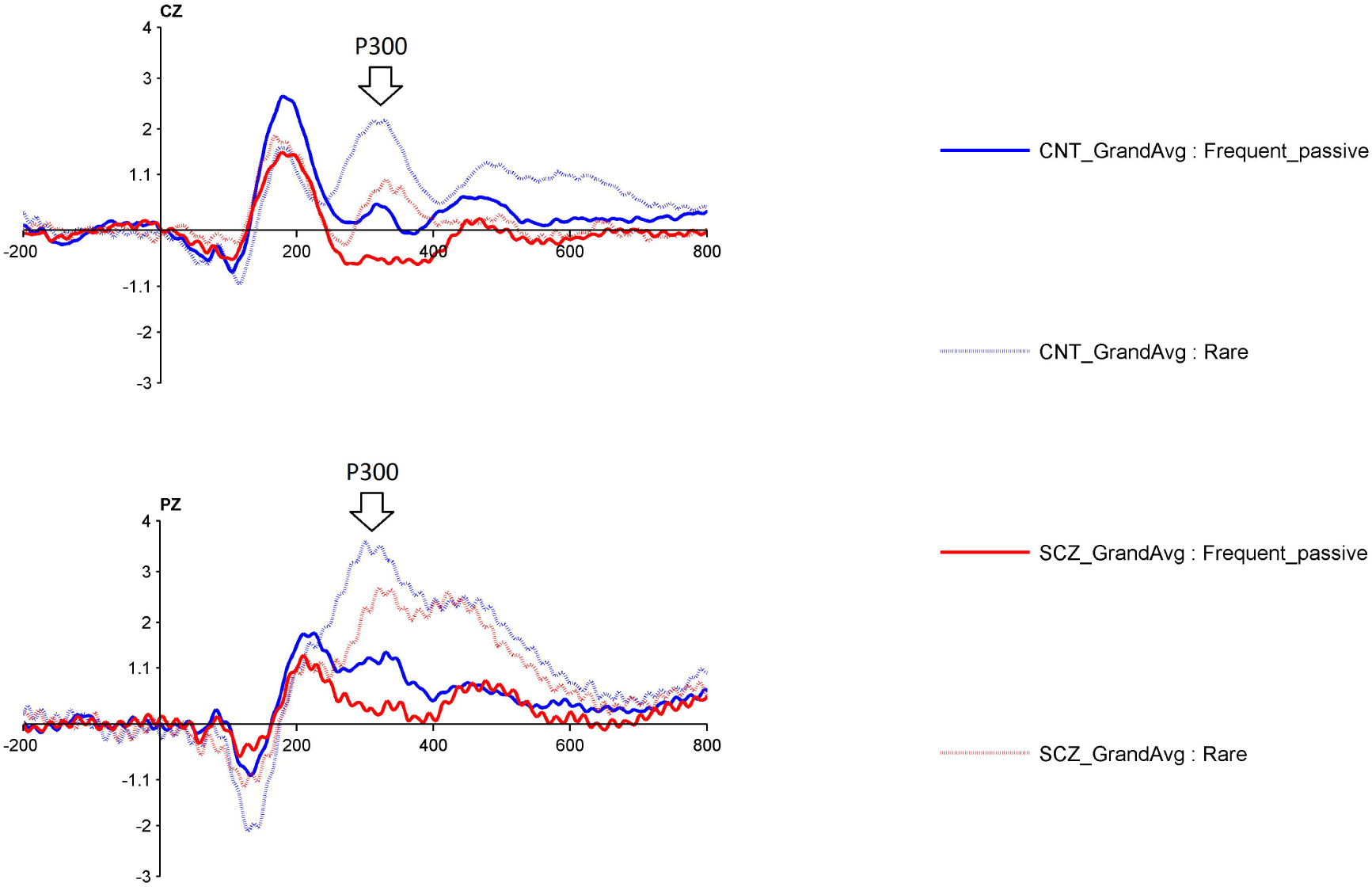
Grand average ERPs showing P300 in patients with schizophrenia (red) and healthy controls (blue). First tones of the standard and deviant tone-pairs were the time-locking stimuli. Arrows denote the ERP component studied.

## Discussion

The current study examined P50 suppression, MMN, ERN and P300 ERPs among SCZ patients, acquired simultaneously during a novel paradigm. We found no significant between group differences for P50 suppression, MMN and P300 ERPs, whereas ERN showed significant reduction in SCZ patients. Additionally, we found a relatively unknown N1-P2 suppression deficit among SCZ patients, during paired tone presentation. To the best of our knowledge, this is the first study involving SCZ patients that reports the above four ERPs evoked simultaneously.

Our results are not in agreement with the earlier studies reporting deficits in P50 (positive ERP component occurring 50ms post-stimulus)(13) as a measure of impaired sensory processing, MMN (mis-match negativity ERP component occurring 100-200ms post-stimulus)(14) which reflects impaired auditory sensory memory, and P300 (positive ERP component occurring ∼300ms post-stimulus)(15) suggesting impaired automatic attention and contextual updating of working memory. Moreover, improvement in MMN and P300 deficits have also been reported to suggest prognosis(16). However the inconsistency between present and prior results could be attributable to methodological differences. Firstly, most (if not all) of the prior ERP studies have used traditional paradigms, where only one or two kinds of stimuli are presented in a session. This is in contrast to a daily life setting, where we typically encounter multiple stimuli of different modalities and context. A recent study has even reported that schizophrenia patients who showed ERP deficits when presented with uni-sensory stimuli, exhibited no difference from healthy counterparts when the same was examined using multi-sensory stimuli(17). Hence, the multi-sensory stimulus presentation achieved during the current study could have allowed the SCZ patients to utilize their intact multisensory integration faculties and generate corresponding ERPs. Being more ecologically valid, the current findings may reflect the true cognitive performance in SCZ patients. Secondly, the current paradigm has many gamification features such as performance feedback, increasing levels of complexity, etc(3). This would allow better subject engagement in comparison to conventional ERP paradigms and may have influenced the ERP generating brain processes.

We however found deficits in ERN as reported in prior studies among SCZ patients(18) as well as among a more broader group of psychotic disorders(19). One possible explanation for the consistency of the ERN finding with some prior reports could be that schizophrenia pathophysiology is currently thought to involve a failure in the brain’s error prediction mechanism(s)(20). Based on supporting neuroimaging findings for this concept, there is a strong view that altered insular network in schizophrenia may be linked to the altered error prediction mechanisms(21). In fact the significant reduction in N1-P2 suppression seen in SCZ patient in the current study, may not be reported before in a paired tone context, but could have a relation to predictive coding. This link is not surprising in light of the N1/P2 suppression deficits reported for self-generated stimuli, among SCZ patients(22–25). In the paired tone condition, the first tone may trigger the prediction mechanism, but results in reduced suppression reflecting the impairment of the same. Thus both the ERN and N1-P2 suppression deficits suggest a strong error prediction abnormality in schizophrenia that may not get corrected despite the multisensory integration.

For the purposes of exploration, the current study limited its analysis on few channels and ERP components that are commonly reported for these ERPs. A more elaborate analysis involving time-frequency measures (both amplitude and phase measures), non-linear measures (entropy and fractal dimensions) and topography-based measures (micro-states) could be used to verify the current findings. Moreover, a larger sample size involving other psychotic illness group should be used in future studies, due to the overlapping nature of these ERPs and to address the scarcity in multisensory and multi-dimensional ERP studies. Furthermore, multivariate classifier analysis using simultaneously acquired ERPs can be useful in segregating high functioning and low functioning individuals from the schizophrenia group, and may thereby help in developing more personalised therapy.

Overall, the current study highlights the importance of using ERPs in a more ecologically valid setting, which can result in better understanding of the pathophysiology of complex mental illnesses like schizophrenia.

## Acknowledgements

We would like to acknowledge the scientific inputs from Mr. Sumit Sharma, Axxonet Technologies Pvt. Ltd., that were critical for the study. We would like to express our most sincere gratitude to the participants who devoted their time and efforts to take part in this study. We thank NIMHANS administration for providing all support to carrying out the study.

